# Universal protection against SARS-CoV-2 viruses by multivalent mRNA vaccine in mice

**DOI:** 10.1101/2023.11.05.565350

**Authors:** Zhengyang Lei, Shiyao Zhai, Xi Yuan, Runming Wang, Yunpeng Wang, Vijay Pandey, Can Yang Zhang, Jiansong Ji, Dongmei Yu, Zhenglin Chen, Sumin Bian, Peiwu Qin

## Abstract

The continual emergence of new severe acute respiratory syndrome coronavirus 2 (SARS-CoV-2) variants challenges available SARS-CoV-2 vaccines for adequate control of outbreaks. Currently, universal vaccines capable of obviating the need for exact strain matching between mRNA vaccines and circulating viruses are absent. In this study, we designed, manufactured, and evaluated a nucleoside-modified lipid nanoparticle mRNA vaccine, aimed for offering broad-spectrum protection against recent SARS-CoV-2 variants. Additionally, the protection efficiency of monovalent, bivalent, quadrivalent, and XBB.1.5 mRNA vaccines was compared with the proposed universal vaccine. The neutralizing antibody activity against wuhan-1, BA.4/5, XBB.1.5, B.1.1.529, BQ.1.1, EG.5.1 and JN.1 was assessed using enzyme-linked immunosorbent assay, rapid fiber-optic biolayer interferometry-based biosensor, and pseudovirus neutralization test. Our results reveal that the proposed multivalent vaccine affords comprehensive protection against previously circulating, current and previously unidentified SARS-CoV-2 strains.

## Introduction

The SARS-CoV-2 pandemic has caused 760 million infections and 6.9 million deaths (https://covid19.who.int). More than 13 billion vaccine doses have been administrated to prevent the virus’s spread. However, the antigens targeted by the majority of these vaccines originate from strains identified early in the pandemic [1, 2]. The time required for vaccine development and the continual emergence of mutant strains have diminished the effectiveness in providing protection against the latest SARS-CoV-2 variants [3, 4].

SARS-CoV-2 invasion is primarily mediated by the binding between spike (S) protein and human angiotensin-converting enzyme-2 (hACE2). The S1 subunit of the S protein contains a receptor binding domain (RBD) responsible for the recognition and binding to the hACE2 [5, 6]. The S protein has been selected as a major target for vaccine development because of its important role in inducing host immune responses and neutralizing antibodies (NAbs) [7-9]. Since the Wuhan-1 strain was identified in late 2019, a continuous stream of SARS-CoV-2 variants such as Alpha (B.1.1.7), Beta (B.1.1.351), Gamma (P.1), and Delta (AY.1) has been identified, with origins in South Africa, Brazil, and India respectively, which shows immune evasion and increased transmissibility [10, 11]. The emergence of the Omicron (B.1.1.529) variant in November 2021 marked a new phase of variant evolution, with subsequent identification of evolving strains including BA.1, BA.2, BA.4, BA.5, BQ.1, XBB.1.5, EG.5, and JN.1, each contributing to the complex landscape of COVID-19 challenges [12-15]. The accumulated mutation site in the spike protein of the Omicron variants facilitates evasion of antibody recognition [3, 16, 17]. To enhance the vaccine efficacy, bivalent and quadrivalent vaccines have been designed and evaluated [18-21]. The bivalent vaccine conferred protection in mice against previously circulating SARS-CoV-2 including BA.1 and BA.4/5 strains [18]. The quadrivalent vaccine was confirmed for its protection against B.1.351 variant [21]. Additionally, the RBD trimer mRNA vaccine demonstrated protective effects against B.1.351 and B.1.617.2 variants [22]. Nevertheless, the efficacy of these vaccines against the newly emerged strains such as XBB.1.5, EG.5.1 and JN.1 has not yet been evaluated due to the rapid evolution of SARS-CoV-2 [23]. Furthermore, Omicron variants CH.1.1 and CA.3.1, identified prior to EG.5.1, have shown nearly complete resistance to neutralization by bivalent boosters [24]. Therefore, there is an urgent need for a universal SARS-CoV-2 vaccine capable of combating the latest strains, alongside a reassessment of the current vaccines’ protective efficiency against these new variants.

Multivalent vaccines have been developed to protect against the Influenza Virus and Human Papillomavirus, which successfully elicited high levels of cross-reactive and subtype-specific antibodies [1, 25, 26]. Increasing evidence suggests that a multivalent approach to SARS-CoV-2 vaccination could address the loss in efficacy of the current vaccine against continually emerging Omicron variants. Furthermore, the mRNA vaccine has exhibited considerable efficacy, safety, and cost-effectiveness in its manufacturing process [27, 28]. Various materials have been developed for delivering mRNA, including polymers, lipids, and protein derivatives [29]. Especially, Lipid nanoparticles (LNP) have been extensively studied and have demonstrated successful clinical applications in mRNA delivery [30]. The benefits of mRNA vaccines facilitate the development of a multivalent vaccine, ensuring a more straightforward, rapid, and cost-efficient production process. The Moderna LNP formulation was adopted in this study considering its superior efficacy in intramuscular delivery of mRNA and antibody production [31-33]. Consequently, a novel multivalent LNP-mRNA SARS-CoV-2 vaccine was designed and manufactured, followed by careful evaluation of its protection efficacy in mice.

In this research, we designed a nucleoside-modified multivalent mRNA vaccine. Hexa-proline and two alanine substitutions were introduced into each variant of concerns (VOCs), variant of interests (VOIs), and wuhan-1 strain [34, 35]. This twenty-valent vaccine encompasses a wide range of strains including wuhan-1, Alpha (B.1.1.7), Beta (B.1.351), Gamma (P.1), Eta (B.1.525), Lota (B.1.526), Kappa, (B.1.617.1), Epsilon (B.1.427), Epsilon (B.1.429), Delta ((B.1.617.2), Delta (AY.1), and Omicron (B.1.1.529, BA.1, BA.1.1, BA.2.7.5, BA.2.12.1, BA.4.6, BA.4/5, XBB.1, and XBB.1.5). This selection effectively represents the most significant VOCs and VOIs that have been previously widespread or are still circulating. After the characterization of the physicochemical properties of the produced LNP-mRNA, mRNAs were *in vitro* transcribed and quantified to create a cocktail of mRNA mix with equal molar concentration for individual mRNA specie. BALB/c mice were used as experimental animals to evaluate the efficacy of universal vaccine. We evaluated in mice the NAbs responses against multiple strains including Wuhan-1, B.1.1.529, BA.4/5, BQ.1.1, XBB.1.5, EG.5.1, and JN.1. Besides, the efficacy of mRNA vaccines including mRNA-1273 (Wuhan-1), mRNA-1273.214 (Wuhan-1/BA.1), mRNA-XBB.1.5, and mRNA-quadrivalent (Alpha/Beta/Delta/BA.1) were reassessed for comparison. The activity of NAbs in immunized mice was assessed through various methods including the enzyme-linked immunosorbent assay (ELISA), a rapid fiber-optic biolayer interferometry-based (FO-BLI) biosensor and pseudovirus neutralization test, which minimizes the assessment bias of single assay [36, 37]. The results revealed that monovalent, bivalent, and quadrivalent vaccines failed to provide sufficient protection against XBB.1.5 and BQ.1.1 strains. In contrast, the multivalent vaccine can provide stronger protection against the latest JN.1 strain where EG.5.1. BQ.1.1, and JN.1 strain is not included in our universal vaccine design. Meanwhile, the multivalent vaccine we proposed demonstrated comprehensive protection against both previous and current SARS-CoV-2 variants, laying a foundation for the development of next-generation universal vaccines against SARS-CoV-2 variants.

## Results

As illustrated in **Figure 1**, we synthesized the mRNA and prepared LNP-RNA vaccines using microfluidic technology. The prepared vaccine was administered to BALB/c mice, with the initial injection on day 0, followed by blood collection on day 21, and a second injection on the same day. Blood was collected again 14 days after the second injection. Serum was extracted from the whole blood samples for the evaluation of neutralizing antibodies. To comprehensively assess the activity of the NAbs, three different methods were employed: a commercialized ELISA kit, the developed FO-BLI biosensor, and a Vesicular Stomatitis Virus (VSV) pseudovirus neutralization test. The bottom section of the figure details the specific virus strains covered by our multivalent vaccine and their periods of prevalence.

**Figure 1.**
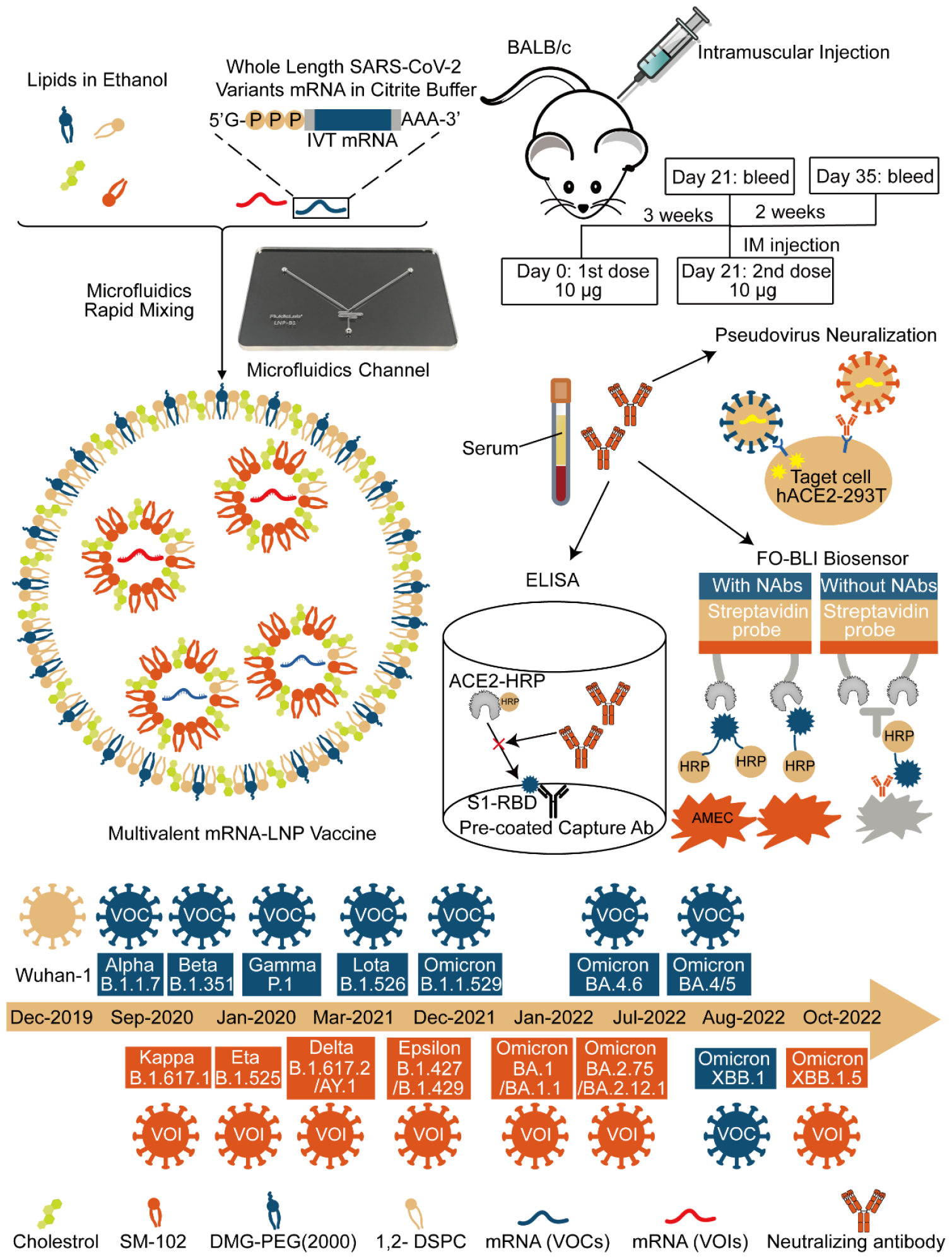
Schematic of the multivalent mRNA vaccine production and its protection efficacy evaluation. 5’-cap mRNA sequences from VOCs and VOIs were synthesized through in vitro transcription and a two-step enzymatic reaction. VOCs: variants of concerns, VOIs: variants of interests. To prepare the mRNA-LNP formulation, four types of lipids, previously dissolved in ethanol, were rapidly mixed with mRNA solution by microfluidics, which was dissolved in citrate buffer. The mRNA-LNP vaccine was then administrated to BALB/C mice via intramuscular injection. Serum samples were collected from immunized mice after the first and booster vaccinations to assess the vaccine’s effectiveness. At the bottom, we illustrated the strains included in the twenty-valent vaccine and the timelines for their classification as VOCs or VOIs. The NAbs activity against SARS-CoV-2 variants was evaluated by ELISA, FO-BLI biosensor, and pseudovirus neutralization test. DSPC: 1,2-Distearoyl-sn-glycero-3 phosphocholine, DMG-PEG2000: 1-monomethoxypolyethyleneglycol-2,3-dimyristylglycerol with polyethylene glycol of average molecular weight 2000. FO-BLI: fiber-optic biolayer interferometry-based biosensor. AMEC: 3-Amino-9-ehtylcarbazole.

The administration of bivalent, quadrivalent and XBB.1.5 vaccines, incorporating targeting antigens from variants such as the Wuhan-1, Alpha, Beta, Gamma, Omicron, and XBB.1.5 spike could enhance efficacy against SARS-CoV-2 variants. This study aims to evaluate the protective efficacy of a proposed multivalent mRNA vaccine protection efficacy and to compare it with that of currently used clinical mRNA vaccines. We developed a total of five LNP-encapsulated mRNA vaccines: mRNA-1273 (wuhan-1), mRNA-1273.214 (wuhan-1/BA.1), mRNA-XBB.1.5, mRNA-quadrivalent (Alpha/Beta/Delta (AY.1) /BA.1), and mRNA-multivalent (twenty-valent). The mRNA constructs design including 5’ untranslated region (5’-UTR), protein coding sequence (CDS), and 3’-UTR was detailed **(Figure 2A)**. The 5’-UTR derived from human α-globin gene was used due to its superior translation efficiency [38]. Hexa-proline and alanine substitutions were introduced to the spike coding sequence of each SARS-CoV-2 strain. The 3’-UTR was a combination of the amino-terminal enhancer (AES) and the human mitochondrial 12S rRNA (mtRNR1), which contributed to increasing protein expression and mRNA stability [39]. These sequences were followed by a 30-mer poly (A) tail, a 10-nucleotide linker sequence (GCAUAUGACU), and 70 adenosine residues, which enhanced protein expression and could be in vitro transcribed from the DNA template without poly (A) addition [40]. Besides, the optimization of the CDS is aligned with that of Moderna-1273, with additional hexa-proline and two alanine substitutions introduced to the Wuhan-1, VOCs, and VOIs mRNA structures. The LNP formulation and production of LNP-mRNA vaccine was adopted as previously described **(Figure 2B)** [22]. Gel electrophoresis was used to evaluate mRNA integrity **(Figure 2C)**. The physiochemical properties of each mRNA-LNP vaccine, including the size, polydispersity index (PDI), and zeta potential were investigated **(Figure 2D and Figure S1A)**. The results showed that the particle sizes of the five LNP-mRNA vaccines we developed are approximately 100 nm, showing excellent uniformity with a PDI of no more than 0.2. The zeta potential and encapsulation rate of the manufactured LNP-mRNA vaccines showed a high level of consistency, which guaranteed uniform antigen expression efficacy [41]. The uniformity of the particle size distribution and LNP morphology were also investigated by a transmission electron microscope (TEM) **(Figure S1B)**.

**Figure 2.**
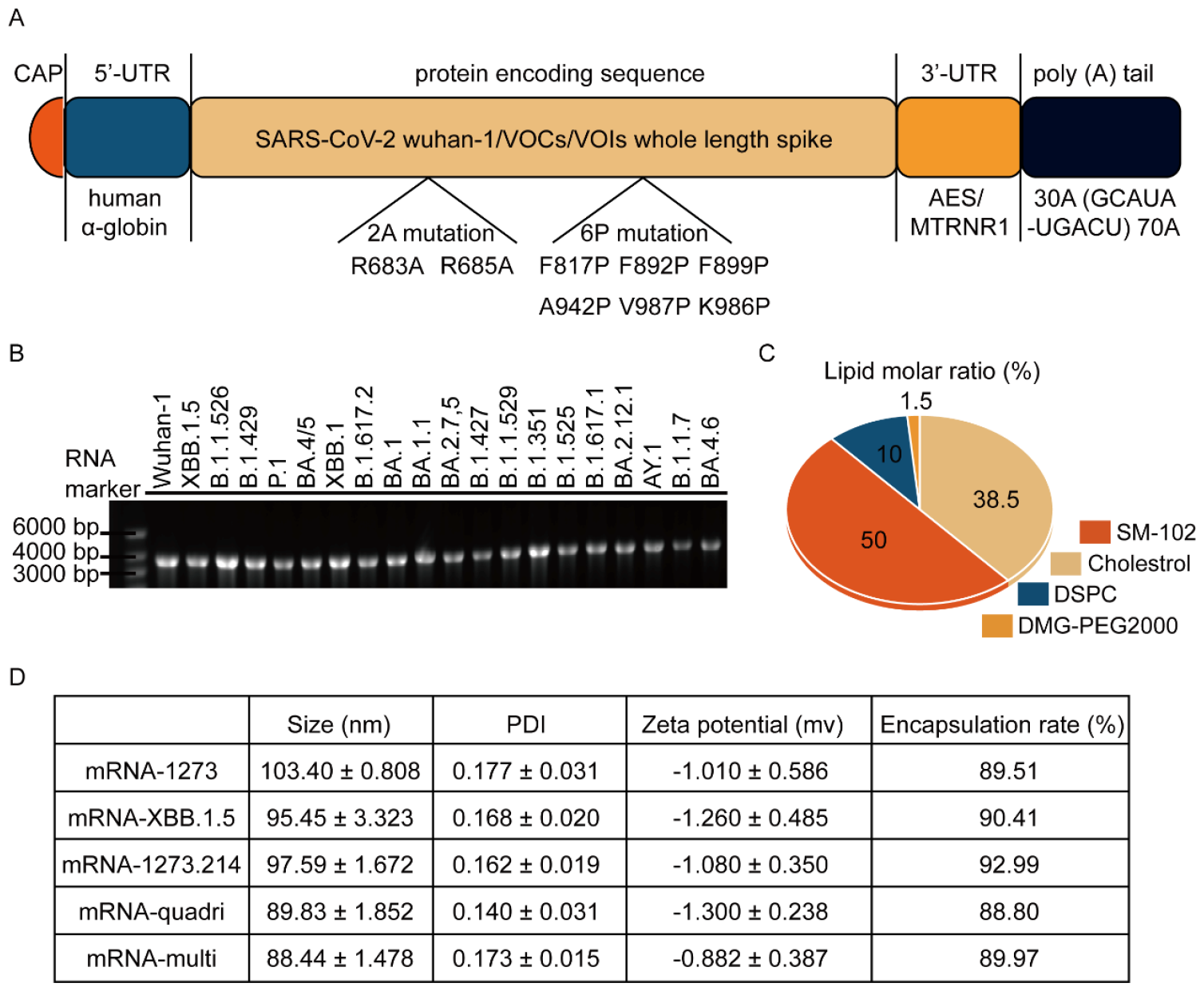
Physicochemical characterization of mRNA-LNPs. (A) Schematic representation of mRNA constructs. (B) Results of agarose gel electrophoresis for in vitro transcribed mRNA products. (C) A pie chart showing the molar ratio of lipids for LNP formulation. DSPC: 1,2-Distearoyl-sn-glycero-3 phosphocholine, DMG-PEG2000: 1-monomethoxypolyethyleneglycol-2,3-dimyristylglycerol with polyethylene glycol of average molecular weight 2000. (D) A table detailing the physicochemical properties of the produced mRNA-LNP vaccines.

As described in **Figure 3A**, we initially vaccinated BALB/c mice with a 10 μg total dose of each vaccine, administrating the second does after three weeks **(Figure 3A)**. The inhibitory capabilities of serum antibodies against the XBB.1.5 strain were evaluated using an RBD ELISA kit three weeks after the first dose (day 21) and two weeks after the second dose (day 35). ELISA results on day 21 indicated the highest NAbs inhibitory activity in sera from mice vaccinated with mRNA-XBB.1.5, followed by the mRNA-multivalent group. The other groups, including those vaccinated with mRNA-1273, mRNA-1273.214, and mRNA-quadrivalent elicited significantly low neutralizing antibodies responses **(Figure 3B)**. Furthermore, ELISA results on day 35 showed that a booster dose of XBB.1.5 and multivalent vaccines enhanced the inhibition rates, nearing 100%. Meanwhile, those receiving monovalent, bivalent, and quadrivalent vaccines showed a slight improvement. Next, we evaluated the protective efficacy of the various vaccines we developed against the wuhan-1 strain and early identified Omicron (B.1.1.529). The results showed that, even with serum diluted a hundredfold, the inhibition efficiency of the five vaccines remained high. However, the XBB.1.5 group’s inhibition rate was lower. Additionally, the bivalent, quadrivalent, and multivalent groups maintained high inhibition rates against the B.1.1.529 strain, while the wuhan-1 and XBB.1.5 group showed reduction in their inhibitory effectiveness **(Figure 3C)**. Subsequent evaluations of protection efficacy against other previously dominant SARS-CoV-2 strains including BA.4/5, and BQ.1.1, revealed that multivalent and XBB.1.5 vaccines were more effective in inhibiting these two strains. Besides, the inhibitory efficiency of the wuhan-1, bivalent, and quadrivalent vaccines against BA.4/5 and BQ.1.1 were significantly lower than that of the multivalent vaccines **(Figure 3D)**. To further validate our results, we employed the FO-BLI biosensor to detect the specific RBD NAbs in the serum of vaccinated mice, with the underlying principle illustrated **(Figure S2A)**. We found that when the serum is diluted a hundred-fold, the inhibition rate against BA.4/5 and BQ.1.1 were about 80%, slightly lower than the inhibition rate detected by ELISA in a ten-fold dilution. Besides, we confirmed that at a ten-fold dilution of serum, the inhibition rate against XBB.1.5 strain remains consistent with the ELISA results **(Figure S2B)**, which further validated our results and demonstrated the accuracy of the developed biosensor. Next, we observed the variations in the inhibition rate against these several virus strains when the serum is gradiently diluted. The serum samples from mice with vaccinated mRNA-XBB.1.5 and mRNA-multivalent were mixed to evaluate the inhibitory activity of the NAbs. We found that when the mixed sample was diluted a ten-fold, the inhibitory rate against XBB.1.5 is consistent with single sample results, and when diluted a hundred-fold, the inhibitory activity significantly decreased, approaching zero **(Figure S2C)**. Meanwhile, when the mixed sample was diluted a hundred-fold, the inhibition rate against BA.4/5 and BQ.1.1 was consistent with the results from individual sample testing, and when diluted to nearly a thousand-fold, the inhibition rate still remained around 30%.

**Figure 3.**
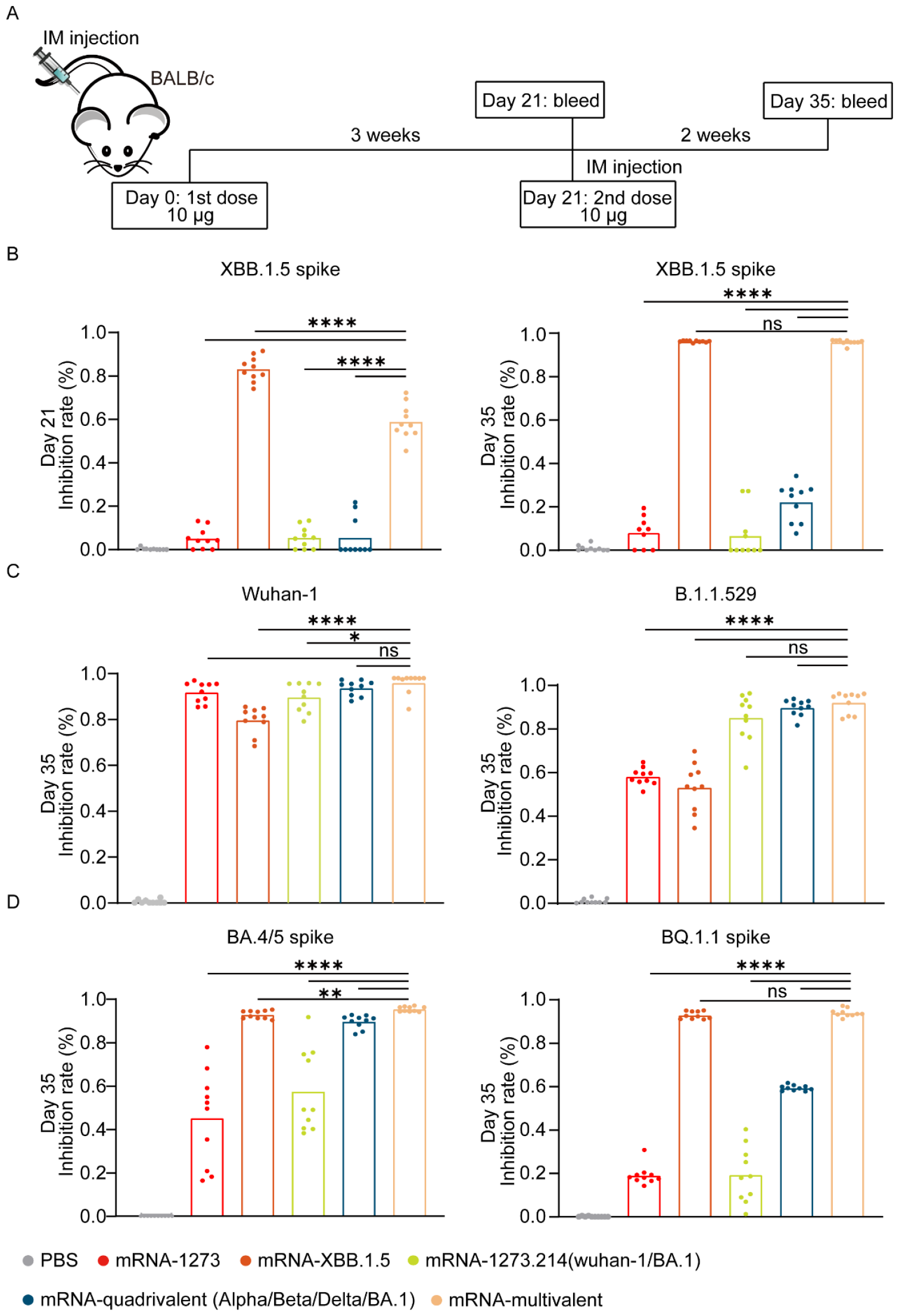
Serum anti-RBD neutralizing antibodies responses in BALB/c evaluated by ELISA. (A) Six-to-eight-week-old BALB/c mice were immunized over a 3-week interval with PBS or 10 μg dose of mRNA vaccine in total. Mice serum was collected at 3 weeks following the initial vaccine administration and 2 weeks following the second boost vaccination. (B) Serum anti-RBD neutralizing antibody activity against XBB.1.5 strain evaluated by ELISA at day 21 and day 35 (n = 10 mice per group in per experiment). The serum samples were diluted tenfold (C) Serum anti-RBD neutralizing antibody activity against wuhan-1 and B.1.1.529 evaluated by ELISA at day 35. The serum samples were diluted a hundred-fold (D) Serum anti-RBD neutralizing antibody activity against BA.4/5 and BQ.1.1 evaluated by ELISA at day 35. The serum samples were diluted ten-fold. Statistical significance was calculated using an unpaired Student’s t test (two-tailed), \**p* < 0.05, \*\**p* < 0.01, \*\*\**p* < 0.001, \*\*\*\**p* < 0.001.

The ELISA-based test and FO-BLI biosensor only detect the antibodies that inhibit the RBD/ACE2 interaction. However, they do not identify non-RBD neutralizing antibodies that may also neutralize the virus. Therefore, A VSV-based neutralization assay, with pseudoviruses displaying spike proteins of the wuhan-1, B.1.1.529, XBB.1.5, BQ.1.1, EG.5.1, and the latest JN.1 was conducted to confirm the findings. Besides, we performed gradient dilutions on serum samples to obtain accurate serum NAbs titers and the half maximal effective concentration (EC_50_) was calculated as described [42]. As a result, low levels of NAbs against XBB.1.5 were induced by mRNA-1273 (GMT: 30), mRNA-1273.214 (GMT: 30), and mRNA-quadrivalent (GMT: 30). The mRNA-XBB.1.5 (GMT: 5196) and mRNA-multivalent (GMT: 2201) showed a strong inhibitory response against XBB.1.5 **(Figure 4A)**. Next, we found that the five vaccines all provided strong protection against the wuhan-1 strain (GMT>7290). Additionally, against the B.1.1.529 virus strain, the quadrivalent (GMT: 1439) and multivalent vaccines (GMT: 3922) elicited higher NAbs responses. While the mRNA-wuhan-1 (GMT: 611), mRNA-XBB.1.5 (GMT: 433), and mRNA-bivalent (GMT: 977) induced lower NAbs responses. Besides, mRNA-XBB.1.5 (GMT: 4524) and mRNA-multivalent (GMT: 2118) both induced strong NAbs activity against EG.5.1. Furthermore, against the BQ.1.1 virus strain, we found that both of vaccines were capable of eliciting strong NAbs activity, with the mRNA-multivalent (GMT: 6501) response being significantly higher than that of the mRNA-XBB.1.5 (GMT: 1447). Finally, in our evaluation against the most recently circulating JN.1 virus strain, the results showed that mRNA-XBB.1.5 (GMT: 131) exhibited an extreme reduction. Meanwhile, the NAbs activity induced by the mRNA-multivalent (GMT: 875) was notably higher.

**Figure 4.**
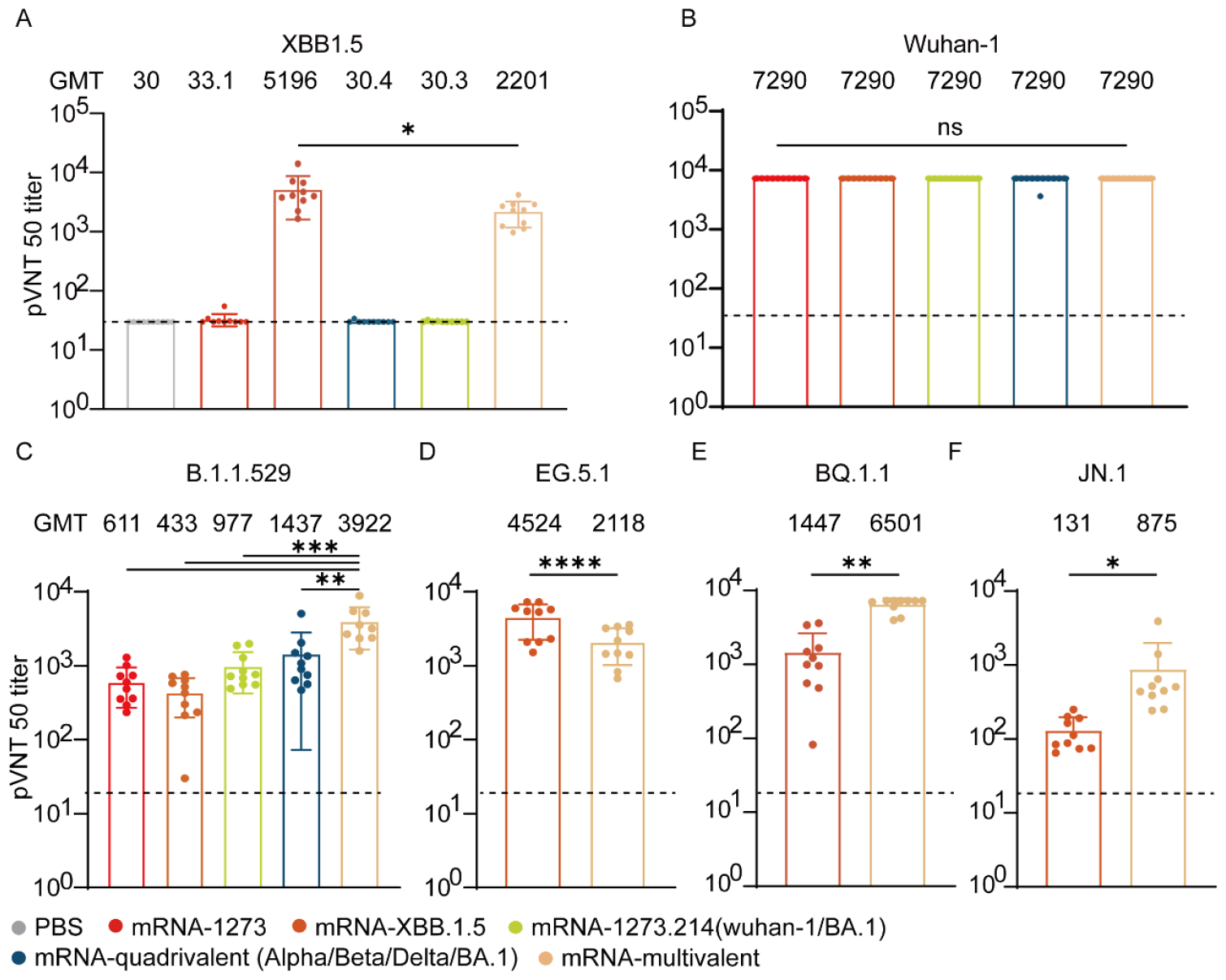
Neutralizing antibodies responses in BALB/c mice after immunization with multivalent vaccine and state-of-the-art vaccines. (A) Neutralizing activity of serum vaccinated with five different mRNA vaccines at day 35 against VSV pseudoviruses displaying the spike proteins of XBB.1.5. (n=10 per mice group, one experiment; tops of boxes show the geometric mean of neutralizing antibody titer (GMT), which are indicated above each column, and dotted lines show LOD). LOD: limit of detection. (B-C) Neutralizing activity of serum vaccinated with five different mRNA vaccines at day 35 against VSV pseudoviruses displaying the spike proteins of wuhan-1 and B.1.1.529. (D-F) Neutralizing activity of serum with two promising vaccines at day 35 against VSV pseudoviruses displaying the spike proteins of EG.5.1, BQ.1.1, and JN.1.

## Discussion

To evaluate the protection efficacy of the proposed universal mRNA vaccine, multiple methods were utilized to evaluate the NAbs activity against multiple SARS-CoV-2 strains. In the ELISA evaluations for the activity of anti-RBD NAbs against XBB.1.5 strains, we noted that post-first immunization, mRNA-XBB.1.5 and mRNA-multivalent significantly outperformed the inhibition rates than other groups. Then, following the administration of the booster dose, the inhibition rate of both vaccines achieved nearly 100%. Moreover, after the administration of the second booster, the mRNA-quadrivalent showed a slight increase in inhibition rate compared to the first dose, while the inhibition rate of the mRNA-1273 and mRNA-bivalent remained low. These data demonstrated that superior protective efficacy when the mRNA vaccine sequence aligned with the virus strain. Besides, mRNA-multivalent had the potential to induce RBD NAbs activity comparable to that triggered by mRNA-XBB.1.5 **(Figure 3B)**. To further assess the efficacy of multivalent vaccine, we evaluated the anti-RBD NAbs against various strains including Wuhan-1, B.1.1.529, BA.4/5, and BQ.1.1. The results indicated that high inhibition rate against Wuhan-1 strain in all five strains, with XBB.1.5 showing lower inhibition capability compared to the others, reinforcing the critical role of antigen-virus sequence matching **(Figure 3C)**. Against the B.1.1.529 strain and the other two variants, the mRNA-multivalent maintained high inhibition rates. Notably, as the virus evolved, these strains emerged sequentially, demonstrating a gradual decrease in inhibition efficiency for mRNA-1273, mRNA-bivalent, and mRNA-quadrivalent, highlighting the challenges posed by viral mutations and the potential benefits of multivalent strategies in maintaining broad protective efficacy **(Figure 3D)**. Next, we utilized the previously developed biosensor to evaluate the RBD NAbs of the promising mRNA-XBB.1.5 and mRNA-multivalent. The single sample inhibition results were consistent with those obtained from the ELISA results, which confirmed the reliability of our data. Due to the difficulty in determining which vaccines was better based on the ten-fold and hundred-fold dilutions, the mixed serum sample was diluted gradiently to evaluate these two promising vaccines efficacy **(Figure S2C)**. Our results showed that the inhibitory efficiency of both vaccines decreases as the dilution factor increases, and at the highest dilution factor, their inhibitory efficiencies are comparable. However, it’s noteworthy that both vaccines exhibited higher inhibitory effects against the BA.4/5 and BQ.1.1 strains compared to their inhibitory action against the XBB.1.5 strain.

To further evaluate the efficacy of various vaccines, including the N-terminal domain (NTD) NAbs [43], we conducted a VSV pseudovirus neutralization tests, expressing a full length XBB.1.5, wuhan-1, B.1.1.529, EG.5.1, BQ.1.1, and JN.1 spike, with gradiently diluted samples. Due to the limitations of dilution factors, we did not obtain precise PVNT50 titer values for each vaccine against the Wuhan-1 virus strain. However, the results showed that their GMT were all greater than 7290, confirming the efficacy of several vaccines against the original strain **(Figure 4B)**. Against XBB.1.5 virus strain, mRNA-XBB.1.5 demonstrated superior protective efficacy compared to multivalent vaccines, evidenced by higher GMT values. Additionally, against B.1.1.529 strain, mRNA-XBB.1.5 and mRNA-1273 showed significantly lower protection efficacy, with mRNA-bivalent, mRNA-quadrivalent, and mRNA-multivalent vaccines displaying a gradual increase in efficacy. This underscores that vaccine efficacy is highest when the sequence of vaccine matches the virus strain. Furthermore, our designed multivalent vaccine maintains robust protection against multiple virus strains **(Figure 4A−4C)**.

Based on our results, the mRNA-XBB.1.5 and mRNA-multivalent vaccines are identified as the most promising candidates against strains emerging after XBB.1.5. Consequently, we evaluated their protective efficacy against several latest virus strains, including EG.5.1, BQ.1.1, and JN.1, which their sequences were not covered in the mRNA-multivalent **(Figure 4D−4F)**. Our data indicated that the mRNA-multivalent vaccine’s protective efficacy against EG.5.1 is lower than that of the mRNA-XBB.1.5. Meanwhile, against BQ.1.1 virus strain, it significantly outperformed the mRNA-XBB.1.5, which demonstrated good GMT level against these two recent virus strains. Notably, the mRNA-multivalent vaccine’s efficacy against JN.1 was significantly higher than that of the mRNA-Xbb.1.5, showcasing its broad-spectrum protective capabilities and highlighting the advantage of multivalent formulations in adapting to the evolving virus landscape. Our findings indicated that mRNA-multivalent vaccine provided superior protection against the not only previously circulating virus strains such as wuhan-1, B.1.1.529, and BA.4/5, also recent XBB.1.5, EG.5.1 variants and the latest JN.1. The discovery carries significant implications for vaccine administration, serving as a valuable reference for both commercial production and clinical application. The emergence of EG.5.1 and JN.1 variants occurred after the initiation of this study, and as a result, no variant-specific vaccine targeting EG.5.1 or JN.1 was developed. Nonetheless, both the mRNA-XBB.1.5 and mRNA-multivalent demonstrated remarkable protection efficacy against EG.5.1 variant **(Figure 3D)**. This aligns with the latest findings from Moderna [44]. In contrast to the recurrent seasonal influenza viruses, the mutation and evolution of SARS-CoV-2 tends to be more unpredictable [45, 46]. While rapid and large-scale SARS-CoV-2 genome sequencing can aid in monitoring its evolution, it necessitates enormous resources in terms of materials and finances [47, 48].

In the event of a virus outbreak similar to those we’ve seen in the past, developing a multivalent vaccine at an appropriate time could be key to rapidly halting the spread of the disease and the virus’s evolution. Multivalent vaccines, by targeting multiple strains or variants simultaneously, offer a broader scope of protection compared to monovalent vaccines, which are designed to protect against a single strain. This broader protection can be crucial in the early stages of an outbreak, where the virus might mutate quickly, and time is of the essence to prevent widespread transmission. By preparing and deploying a multivalent vaccine, public health strategies can gain a significant advantage in controlling and eventually eradicating the viral threat. Additionally, the production of mRNA vaccine is notably straightforward and mature, facilitating the swift creation of multivalent vaccine against a virus. Therefore, this proposed universal mRNA-multivalent is poised to play a pivotal role in preventing the COVID-19 pandemic and a multivalent vaccine strategy could be used for preventing potential virus disease pandemic.

## Conclusion

In this study, we introduced an innovative twenty-valent SARS-CoV-2 mRNA vaccine. Our findings conclusively revealed that the mRNA-multivalent vaccine elicited potent, universal neutralizing activity against a spectrum of variants, including wuhan-1, BA.4/5, BQ.1.1, XBB.1.5, EG.5.1, and JN.1 variants. We firmly believe this novel mRNA-multivalent vaccine can establish a valuable foundation and provide critical insight for the development of a universal SAR-CoV-2 vaccine.

## Materials and Methods

### Mice

BALB/c mice aged 6-8 weeks were purchased from Bestest Biotechnology Co., Ltd and subsequently housed at TOP Biotechnology Co., Ltd. Animal experiments were carried out in compliance with approval from the Animal Care and Use Committee of the TOP Biotechnology Co., Ltd.

### Construction of expression vectors encoding the wuhan-1, VOCs and VOIs spike of SARS-CoV-2

Condon-optimized full-length SARS-CoV-2 wuhan-1, VOCs, and VOIs spike were constructed and verified by sequencing, then cloned into a pSLQ1651-sgTelomere (F+E) vector. Proline substitutions (F817P, A892P, A899P, A942P, K986P, and V987P) and alanine substitutions (R683A and R685A) were introduced to each expressing vector to stabilize the trimeric prefusion state of SARS-CoV-2 S protein and enhance immunogenicity. Besides, spike genes contained the following mutations: (a)

B.1.1.7: N503Y, A572D, D616G, P683H, T718I, S984A, D1120H, Δ69-70, and Δ144; (b) 1.3.5.1: L18F, D80A, D217G, R248I, K419N, E486K, N503Y, D616G, and A703V; (c) P.1: L18F, T20N, P26S, D138Y, R190S, K419T, E486K, N503Y, D616G, H657Y, T1029I, and V1178F; (d) B.1.525: Q52R, A67V, E486K, D616G, Q679H, F890L, Δ69-70, and Δ144; (e) B.1.526: L5F, T95I, D255G, E486K, D616G, A703V, and F890P; (f) B.1.617.1: T95I, G142D, E154K, L454R, E486Q, D616G, P683R, and Q1073H; (g) B.1.427: L454R, and D616G; (h) B.1.429: C131I, W152C, L454R, and D616G; (i) B.1.617.2 : T19R, G142D, E156G, L454R, T480K, D616G, P683R, D952N, and Δ157-158; (j) AY.1: T19R, V70F, A224V, W260L, K419N, L454R, T480K, D616G, P683R, D952N, and Δ157-158; (k): B.1.1.529: A67V, T95I, G142D, N211I, L212V, V213R, Δ214E, Δ215P, R216E, G341D, S373L, S375P, S377F, K419N, N442K, G448S, S479N, T480K, E486A, Q495R, G498S, Q500R, N503Y, Y507H, T549K, D616G, H657Y, N681K, P683H, N766K, D798Y, N858K, Q956H, N971K, L983F, Δ69-70, and Δ143-145; (l) BA.2.12.1: T19I, A27S, G142D, V213G, G341D, S373F, S375P, S377F, T378A, D407N, R410S, K419N, N442K, L454Q, S479N, T480K, E486A, Q495R, Q500R, N503Y, Y507H, D616G, H657Y, N681K, P683H, A685G, A687G, S706L, N766K, D798Y, Q954H, N971K, and Δ24-26; (m) BA.1.1: A67V, T95I, G142D, N211I, L212V, V213R, Δ214E, Δ215P, R216E, G341D, R348K, S373L, S375P, S377F, K419N, N442K, G448S, S479N, T480K, E486A, Q495R, G498S, Q500R, N503Y, Y507H, T549K, D616G, H657Y, N681K, P683H, N766K, D798Y, N858K, Q956H, N971K, L983F, Δ69-70, and Δ143-145; (n) BA.2.7.5 : T19I, A27S, G142D, K147E, W152R, F157L, I210V, V213G, G259S, G341H, S373F, S375P, S377F, T378A, D407N, R410S, K419N, N442K, G448S, N462K, S479N, T480K, E486A, Q500R, N503Y, Y507H, D616G, H657Y, N681K, P683H, N766K, D798Y, Q956H, N971K, and Δ24-26; (o) BA.4.6: T19I, A27S, G142D, V213G, G341D, R348T, S373F, S375P, S377F, T378A, D407N, R410S, K419N, N442K, L454R, S479N, T480K, E486A, F488V, Q500R, N503Y, Y507H, D616G, H657Y, N660S, N681K, P683H, N766K, D798Y, Q956H, N971K, Δ24-26, and Δ69-70; (p) BA.1: A67V, T95I, G142D, N211I, L212V, V213R, Δ214E, Δ215P, R216E, G341D, S373L, S375P, S377F, K419N, N442K, G448S, S479N, T480K, E486A, Q495R, G498S, Q500R, N503Y, Y507H, T549K, D616G, H657Y, N681K, P683H, N766K, D798Y, N858K, Q956H, N971K, L983F, Δ69-70, Δ143-145, and Δ1276, (q) BA.4/5: T19I, A27S, G142D, V213G, G341D, S373F, S375P, S377F, T378A, D407N, R410S, K419N, N442K, L454R, S479N, T480K, E486A, F488V, Q500R, N503Y, Y507H, D616G, H657Y, N681K, P683H, N766K, D798Y, Q956H, N971K, Δ24-26, and Δ69-70; (r) XBB.1: T19I, A27S, V83A, G142D, H146Q, Q183E, V213E, G341H, R348T, L370I, S373F, S375P, S377F, T378A, D407N, R410S, K419N, N442K, V447P, G448S, N462K, S479N, T480K, E486A, F488S, F492S, Q500R, N503Y, Y507H, D616G, H657Y, N681K, P683H, N766K, D798Y, Q956H, N971K, Δ24-26, and Δ144; (s) XBB.1.5: T19I, A27S, V83A, G142D, H146Q, Q183E, V213E, G341H, R348T, L370I, S373F, S375P, S377F, T378A, D407N, R410S, K419N, N442K, V447P, G448S, N462K, S479N, T480K, E486A, F488S, F492S, Q500R, N503Y, Y507H, D616G, H657Y, N681K, P683H, N766K, D798Y, Q956H, N971K, Δ24-26, and Δ144.

### Design and production of lipid-nanoparticle encapsulated mRNA vaccines

The mRNA was transcribed from the linearized DNA templated using T7 RNA polymerase reaction with complete replacement of uridine by N1m-pseudouridine (Vazyme, DD4202). The transcription products were then purified (New England Biolabs, T2050L) and the cap1 structure was added for efficient translation and evading the cellular innate immune response (New England Biolabs, M2081 and M0366). Lipid-nanoparticle (LNP) formulation of mRNA was performed as previously described. Briefly, lipids were dissolved in absolute ethanol containing an ionizable lipid SM-102, 1, 2-distearoly-sn-glycero-3-phosphocholine (DSPC), cholesterol, and PEG-lipid with a total molar weight 8 mM and the molar ratio of 50:10:38.5:1.5. The lipid mixture was combined with 50 mM citrate buffer (pH 4.0) containing mRNA at an N/P ratio 6:1, a flow rate ratio 3:1, and total flow rate 12 ml/min through a T-mixer. Formulations were then diafiltrated against 30× volume of PBS (pH 7.4) through a tangential-flow filtration (TFF) membrane with 30 KD molecular weight cut-offs. 10% Sucrose was added as a cryoprotectant. The final solution was sterile-filtered passing through a 0.22 mm filter and stored at -20 °C for further use. All formulations were tested for particle size, distribution, RNA concentration and encapsulation.

### Quality control of LNP-mRNA

Encapsulation of mRNA in the LNP was measured using a Quani-it^TM^ RiboGreen^TM^ RNA Assay Kit according to the manufacturer’s instructions (ThermoFisher Scientific, R11491). The size, polydispersity (PDI) and zeta-potential of mRNA-LNP was measured by dynamic light scattering (DLS) on a Zeta sizer Nano ZS (Malvern instruments, UK). LNP-mRNA was diluted 100 times in PBS (pH 7.4) and added to a cuvette. The dispersant (PBS) refractive index (RI), viscosity values, and material RI were 1335, 1.02, and 1.45, respectively. Scattered light was detected at a backscattering angle of 173°. Results were analyzed using software of Zeta sizer V7.13. The morphology of LNP-mRNA was investigated using the FEI Tecnai T12 G2 cryo-TEM. The samples were stained with 2% uranylacetate as previously described [21].

### SARS-CoV-2 Surrogate Virus Neutralization Test (sVNT) kit (ELISA)

To detect the amount and inhibition rate of neutralizing antibodies from the serum of immunized mice against VOCs, the commercialized ELISA kit (Genscript, Cat#L00871) was used. The tests were conducted according to the manufacture’s protocol. Briefly, the kit and serum samples were removed and left at room temperature (20-25 °C) before running the assay at least 30 min. Serum samples were diluted with sample dilution buffer with a volume ratio of 1:9. Standard stocks were serially diluted as described in the protocol to establish the standard calibration curves. Probe RBD (XBB.1.5, BQ.1.1, and BA.4/5) and enzyme conjugate were diluted with assay buffer with a volume ratio of 1:2000. The diluted serum samples and probe RBD working solution were mixed in a volume ratio of 2:1 and incubated at 37 °C for 30 min. 100 μL mixture was transferred to the captured plate, which was then covered by a plate sealer and incubated at 30 °C for 15 min. Then, the plate was washed with 1× wash solution four times. The residual liquids in the wells were totally removed and 100 μL enzyme conjugate working solution was added. The sealed plate was incubated at 37 °C for 15 min and repeat the wash steps. 100 μL TMB solution was added and the plate was incubated in the dark at 20-25 °C for 15 min. 50 μL Stop solution was added to each well. Finally, the absorbance at 450 nm was recorded immediately (Spark Tecan, Switzerland).

### VSV Pseudovirus neutralization assay

The spike gene used in VSV assay contains the following mutation sites: (a) B.1.1.529: A67V, △69, △70, T95I, G142D, △143, △144, △145, △211, L212I, ins214EPE, G339D, S371L, S373P, S375F, K417N, N440K, G446S, S477N, T478K, E484A, Q493R, G496S, Q498R, N501Y, Y505H, T547K, D614G, H655Y, N679K, P681H, N764K, D796Y, N856K, Q954H, N969K, L981F; (b) BQ.1.1: T19I, ΔL24 ΔP25-26, A27S, Δ69-70, G142D, V213G, G339D, R346T, S371F, S373P, S375F, T376A, D405N R408S, K417N, N440K, K444T, L452R, N460K, S477N, T478K, E484A, F486V, Q498R, N501Y, Y505H, D614G, H655Y, N679K, P681H, N764K, D796Y, Q954H, N969K; (c) XBB.1.5: T19I, L24-, P25-, P26-, A27S, V83A, G142D, Y144-, H146Q, Q183E, V213E, G252V, G339H, R346T, L368I, S371F, S373P, S375F, T376A, D405N, R408S, K417N, N440K, V445P, G446S, N460K, S477N, T478K, E484A, F486P, F490S, Q498R, N501Y, Y505H, D614G, H655Y, N679K, P681H, N764K, D796Y, Q954H, N969K; EG.5.1: T19I, L24-, P25-, P26-, A27S, Q52H, V83A, G142D, Y144-, H146Q, Q183E, V213E, G252V, G339H, R346T, L368I, S371F, S373P, S375F, T376A, D405N, R408S, K417N, N440K, V445P, G446S, F456L, N460K, S477N, T478K, E484A, F486P, F490S, Q498R, N501Y, Y505H, D614G, H655Y, N679K, P681H, N764K, D796Y, Q954H, N969K; JN.1: T19I, R21T, L24-, P25-, P26-, A27S, S50L, H69-, V70-, V127F, G142D, Y144-, F157S, R158G, N211-, L212I, V213G, L216F, H245N, A264D, I332V, G339H, K356T, S371F, S373P, S375F, T376A, R403K, D405N, R408S, K417N, N440K, V445H, G446S, N450D, L452W, L455S, N460K, S477N, T478K, N481K, V483-, E484K, F486P, Q498R, N501Y, Y505H, E554K, A570V, D614G, P621S, H655Y, N679K, P681R, N764K, D796Y, S939F, Q954H, N969K, P1143L. ACE2-HEK293T
cells were used as target cells for the neutralization assay and maintained in DMEM supplemented with 10% FBS. To perform the neutralization assay, immunized mice serum samples were heat-inactivated for 45 minutes at 56 °C. The sera at a starting dilution of 1:30 (six dilutions) were threefold diluted with DMEM, and then mixed with an equal volume of pre-titrated pseudotyped virus. After incubation at 37 °C for 2 h, the inoculum or virus-serum mix was subsequently used to infect hACE2-293T cells (3×10^4^ cells/ well) in the 96-well microplate (Corning, Cat^#^3917). After incubation for 48 h, an equal volume of One-Glo reagent (Promega, E6120) was added to the culture medium for readout. The percentage inhibition was calculated based on based on relative light units (RLUs) of the virus control and subsequently analyzed using a four-parameter logistic curve (GraphPad Prism 9.0).

### FO-BLI biosensor detecting assay

The FO-BLI detection protocol is as previously described with some modification [36]. The FO-BLI NAbs biosensor was recently proven to be highly correlated with a pseudovirus neutralization assay in identifying neutralizing capacities towards wide type and Omicro variants [37]. Shortly, Biotinylated hACE2-functionalized sensors were employed to capture SARS-CoV-2 NAbs via competitive binding. AMEC, as an environment- and user-friendly signal enhancer, and HRP-conjugated RBD (RBD-HRP) were utilized in the assay for signal amplification and detection, and competition, respectively. In order to achieve adequate binding for the NAbs biosensor, 5 μg/mL of biotinylated hACE2 in PBS-T buffer was utilized to induce loading shifts of approximately 1.0 nm. Prior to testing, RBD-HRP was combined in equal volumes with spiked samples. Between steps, washing was performed, and the signals produced were proportional to the quantity of RBD-HRP in the underlying complex. Specifically, SARS-CoV-2 (BA.4/BA.5/BA.5.2) Spike RBD Protein, SARS-CoV-2 XBB (Omicron) Spike RBD Protein and SARS-COV-2 BQ.1.1 (Omicron) Spike RBD Protein were ordered from SinoBiological (Beijing, China), while ImmPACT* AMEC Red* Substrate Kit was from Vector Laboratories (Shanghai, China). All the mice sera were diluted 100-fold using a high-salt sample buffer, (i.e., PBS (10 mM, pH 7.4) containing 274 mM NaCl, 0.02% Tween20 and 0.1% BSA) to minimize the non-specific matrix interference. A mixed healthy mice serum collected from a mouse without vaccine administration was utilized as a control to calculate the relative inhibitory rate. The test was performed using a Octet^®^ RED96E 8-Channel System (Sartorius Group, Gottingen, Germany). Determination of neutralizing capacities by the FO-BLI biosensor was completed automatically within ten minutes.

## Author contribution

P.Q., S.B., and Z.C. supervised the study. Z.L. performed the experiment, collected data, analyzed data, and wrote the first draft of the manuscript. S.B. detected NAbs with FO-BLI biosensor, analyzed data, and revised the manuscript. S.Z. and X.Y. helped with the animal experiment. R.W., H.X. K.D., Y.C., P.W., and P.V. and R.W. helped revise the manuscript.

## Acknowledgment

We thank the funding support from the Tsinghua Shenzhen International Graduate School Cross-disciplinary Research and Innovation Fund Research Plan, JC2022009; National Natural Science Foundation of China 31970752, 82104122; Science, Technology, Innovation Commission of Shenzhen Municipality JCYJ20190809180003689, JSGG20200225150707332, JCYJ20220530143014032, ZDSYS20200820165400003, WDZC20200820173710001, WDZC20200821150704001, JSGG20191129110812708; Shenzhen Bay Laboratory Open Funding, SZBL2020090501004; Department of Chemical Engineering-iBHE special cooperation joint fund project, DCE-iBHE-2022-3; and Bureau of Planning, Land and Resources of Shenzhen Municipality, (2022) 207.

## References

1. Final efficacy, immunogenicity, and safety analyses of a nine-valent human papillomavirus vaccine in women aged 16–26 years: a randomised, double-blind trial. The Lancet, 2017.

2. Dong, Y., et al., A systematic review of SARS-CoV-2 vaccine candidates. Signal Transduct Target Ther, 2020. 5(1): p. 237.

3. Cao, Y., et al., Omicron escapes the majority of existing SARS-CoV-2 neutralizing antibodies. Nature, 2022. 602(7898): p. 657–663.

4. Zhao, F., et al., Challenges and developments in universal vaccine design against SARS-CoV-2 variants. NPJ Vaccines, 2022. 7(1): p. 167.

5. Qi, H., et al., The humoral response and antibodies against SARS-CoV-2 infection. Nat Immunol, 2022. 23(7): p. 1008–1020.

6. Bao, L., et al., The pathogenicity of SARS-CoV-2 in hACE2 transgenic mice. Nature, 2020. 583(7818): p. 830–833.

7. Du, L., Y. Yang, and X. Zhang, Neutralizing antibodies for the prevention and treatment of COVID-19. Cell Mol Immunol, 2021. 18(10): p. 2293–2306.

8. Kim, C., et al., A therapeutic neutralizing antibody targeting receptor binding domain of SARS-CoV-2 spike protein. Nat Commun, 2021. 12(1): p. 288.

9. Pierri, C.L., SARS-CoV-2 spike protein: flexibility as a new target for fighting infection. Signal Transduct Target Ther, 2020. 5(1): p. 254.

10. Radvak, P., et al., SARS-CoV-2 B.1.1.7 (alpha) and B.1.351 (beta) variants induce pathogenic patterns in K18-hACE2 transgenic mice distinct from early strains. Nat Commun, 2021. 12(1): p. 6559.

11. Hoffmann, M., et al., SARS-CoV-2 variants B.1.351 and P.1 escape from neutralizing antibodies. Cell, 2021. 184(9): p. 2384–2393 e12.

12. Kumar, S., K. Karuppanan, and G. Subramaniam, Omicron (BA.1) and sub-variants (BA.1.1, BA.2, and BA.3) of SARS-CoV-2 spike infectivity and pathogenicity: A comparative sequence and structural-based computational assessment. J Med Virol, 2022. 94(10): p. 4780–4791.

13. Dejnirattisai, W., et al., SARS-CoV-2 Omicron-B.1.1.529 leads to widespread escape from neutralizing antibody responses. Cell, 2022. 185(3): p. 467–484 e15.

14. Lewnard, J.A., et al., Clinical outcomes associated with SARS-CoV-2 Omicron (B.1.1.529) variant and BA.1/BA.1.1 or BA.2 subvariant infection in Southern California. Nat Med, 2022. 28(9): p. 1933–1943.

15. Yang, S., et al., Fast evolution of SARS-CoV-2 BA.2.86 to JN.1 under heavy immune pressure. Lancet Infect Dis, 2024. 24(2): p. e70–e72.

16. Cao, Y., et al., BA.2.12.1, BA.4 and BA.5 escape antibodies elicited by Omicron infection. Nature, 2022. 608(7923): p. 593–602.

17. Kurhade, C., et al., Low neutralization of SARS-CoV-2 Omicron BA.2.75.2, BQ.1.1 and XBB.1 by parental mRNA vaccine or a BA.5 bivalent booster. Nat Med, 2023. 29(2): p. 344–347.

18. Scheaffer, S.M., et al., Bivalent SARS-CoV-2 mRNA vaccines increase breadth of neutralization and protect against the BA.5 Omicron variant in mice. Nat Med, 2023. 29(1): p. 247–257.

19. Branche, A.R., et al., Comparison of bivalent and monovalent SARS-CoV-2 variant vaccines: the phase 2 randomized open-label COVAIL trial. Nat Med, 2023. 29(9): p. 2334–2346.

20. Izikson, R., et al., Safety and immunogenicity of a high-dose quadrivalent influenza vaccine administered concomitantly with a third dose of the mRNA-1273 SARS-CoV-2 vaccine in adults aged >/=65 years: a phase 2, randomised, open-label study. Lancet Respir Med, 2022. 10(4): p. 392–402.

21. Kang, Y.F., et al., Quadrivalent mosaic HexaPro-bearing nanoparticle vaccine protects against infection of SARS-CoV-2 variants. Nat Commun, 2022. 13(1): p. 2674.

22. Liang, Q., et al., RBD trimer mRNA vaccine elicits broad and protective immune responses against SARS-CoV-2 variants. iScience, 2022. 25(4): p. 104043.

23. Abbasi, J., What to Know About EG.5, the Latest SARS-CoV-2 “Variant of Interest”. JAMA, 2023. 330(10): p. 900–901.

24. Qu, P., et al., Enhanced evasion of neutralizing antibody response by Omicron XBB.1.5, CH.1.1, and CA.3.1 variants. Cell Rep, 2023. 42(5): p. 112443.

25. Arevalo, C.P., et al., A multivalent nucleoside-modified mRNA vaccine against all known influenza virus subtypes. Science, 2022. 378(6622): p. 899–904.

26. Chatterjee, A., The next generation of HPV vaccines: nonavalent vaccine V503 on the horizon. Expert Rev Vaccines, 2014. 13(11): p. 1279–90.

27. Chaudhary, N., D. Weissman, and K.A. Whitehead, mRNA vaccines for infectious diseases: principles, delivery and clinical translation. Nat Rev Drug Discov, 2021. 20(11): p. 817–838.

28. Pardi, N., et al., mRNA vaccines - a new era in vaccinology. Nat Rev Drug Discov, 2018. 17(4): p. 261–279.

29. Hou, X.C., et al., Lipid nanoparticles for mRNA delivery (Aug, 10.1038/s41578-021-00358-0, 2021). Nature Reviews Materials, 2022. 7(1): p. 65–65.

30. Ndeupen, S., et al., The mRNA-LNP platform’s lipid nanoparticle component used in preclinical vaccine studies is highly inflammatory. iScience, 2021. 24(12): p. 103479.

31. Baden, L.R., et al., Efficacy and Safety of the mRNA-1273 SARS-CoV-2 Vaccine. N Engl J Med, 2021. 384(5): p. 403–416.

32. Jackson, L.A., et al., An mRNA Vaccine against SARS-CoV-2-Preliminary Report. N Engl J Med, 2020. 383(20): p. 1920–1931.

33. Pegu, A., et al., Durability of mRNA-1273 vaccine-induced antibodies against SARS-CoV-2 variants. Science, 2021. 373(6561): p. 1372–1377.

34. Hsieh, C.L., et al., Structure-based design of prefusion-stabilized SARS-CoV-2 spikes. Science, 2020. 369(6510): p. 1501–1505.

35. Martinez-Flores, D., et al., SARS-CoV-2 Vaccines Based on the Spike Glycoprotein and Implications of New Viral Variants. Front Immunol, 2021. 12: p. 701501.

36. Bian, S., M. Shang, and M. Sawan, Rapid biosensing SARS-CoV-2 antibodies in vaccinated healthy donors. Biosens Bioelectron, 2022. 204: p. 114054.

37. Bian, S., et al., Dynamic Profiling and Prediction of Antibody Response to SARS-CoV-2 Booster-Inactivated Vaccines by Microsample-Driven Biosensor and Machine Learning. Vaccines, 2024. 12(4): p. 352.

38. Xia, X., Detailed Dissection and Critical Evaluation of the Pfizer/BioNTech and Moderna mRNA Vaccines. Vaccines (Basel), 2021. 9(7).

39. Granados-Riveron, J.T. and G. Aquino-Jarquin, Engineering of the current nucleoside-modified mRNA-LNP vaccines against SARS-CoV-2. Biomed Pharmacother, 2021. 142: p. 111953.

40. Beissert, T., et al., A Trans-amplifying RNA Vaccine Strategy for Induction of Potent Protective Immunity. Mol Ther, 2020. 28(1): p. 119–128.

41. Arteta, M.Y., et al., Successful reprogramming of cellular protein production through mRNA delivered by functionalized lipid nanoparticles. Proceedings of the National Academy of Sciences of the United States of America, 2018. 115(15): p. E3351–E3360.

42. Nie, J., et al., Quantification of SARS-CoV-2 neutralizing antibody by a pseudotyped virus-based assay. Nat Protoc, 2020. 15(11): p. 3699–3715.

43. Chi, X., et al., A neutralizing human antibody binds to the N-terminal domain of the Spike protein of SARS-CoV-2. Science, 2020. 369(6504): p. 650–655.

44. Chalkias, S., et al., Three-month antibody persistence of a bivalent Omicron-containing booster vaccine against COVID-19. Nat Commun, 2023. 14(1): p. 5125.

45. Petrova, V.N. and C.A. Russell, The evolution of seasonal influenza viruses. Nat Rev Microbiol, 2018. 16(1): p. 47–60.

46. Markov, P.V., et al., The evolution of SARS-CoV-2. Nat Rev Microbiol, 2023. 21(6): p. 361–379.

47. Chen, Q., et al., CRISPR-Cas12-based field-deployable system for rapid detection of synthetic DNA sequence of the monkeypox virus genome. J Med Virol, 2023. 95(1): p. e28385.

48. Bahri, M., et al., Laser-Induced graphene electrodes for highly sensitive detection of DNA hybridization via consecutive cytosines (polyC)-DNA-based electrochemical biosensors. Microchemical Journal, 2023. 185.

