## supporting information for "Universal protection against SARS-CoV-2 viruses by multivalent mRNA vaccine in mice"

**Manuscript − Supporting information**

**Title**

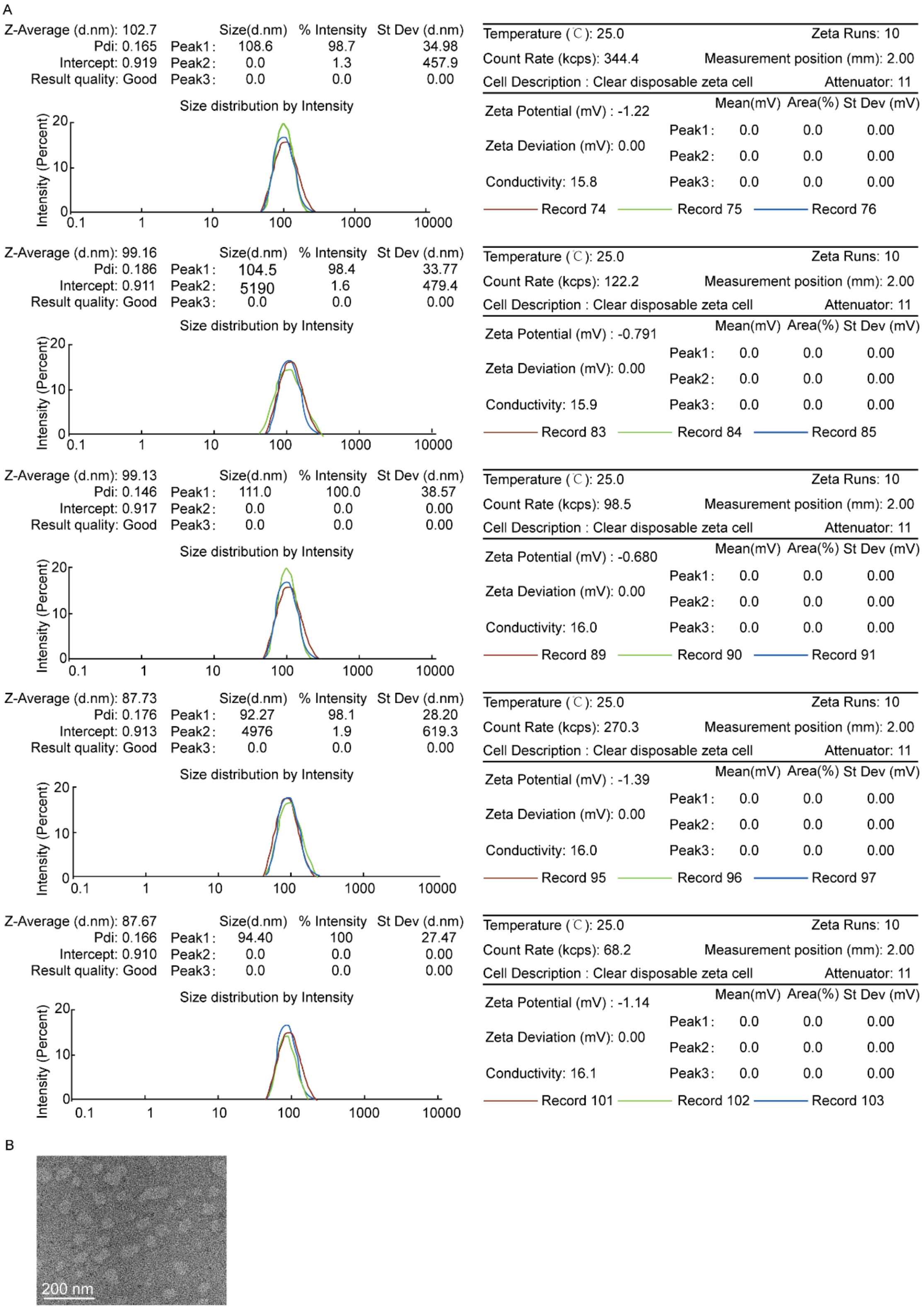


**Figure S1. Characterization of the physicochemical properties of the produced LNP-mRNA vaccines.** (A) The size, PDI, and zeta potential of mRNA-LNP (mRNA-1273, mRNA-XBB.1.5, mRNA-1273.124, mRNA-quadrivalent, mRNA-multivalent) measured by dynamic light scattering (DLS) on a Zeta sizer Nano ZS. (B) A representative image of LNP-mRNA through cryo-TEM imaging, a scale bar of 200 nm is indicated.

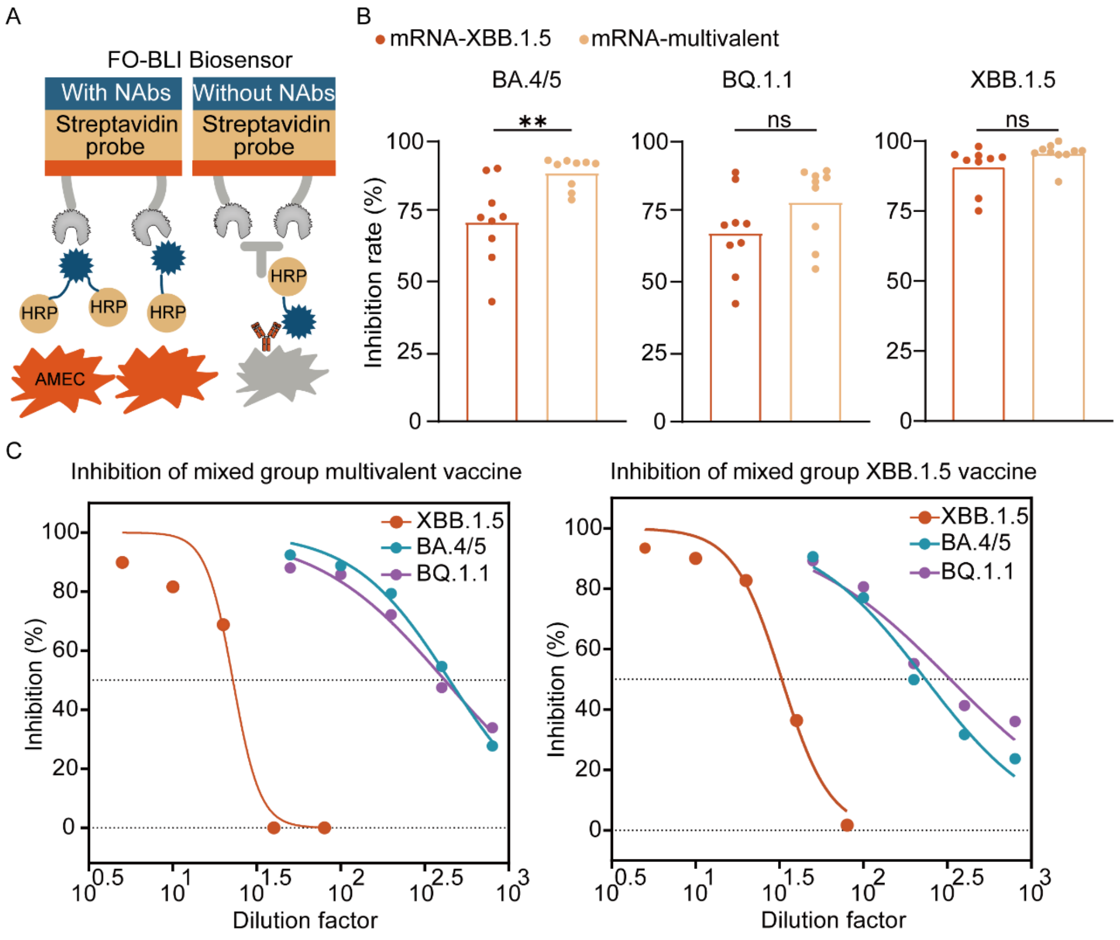


**Figure S2. Neutralizing antibody activity evaluated by FO-BLI biosensor.** (A) Principle of the FO-BLI biosensor for detection of NAbs via a competitive binding format. In the absence of NAbs, the interaction between the HPR-RBD domain of the SARS-CoV-2 spike protein and hACE2 leads to strong detection signals. Conversely, the presence of NAbs inhibits the binding of HRP-RBD to immobilized hACE2 protein. The optical signals were significantly enhanced by the addition of DAB in both bioassays. (B) Serum neutralizing antibody activity against BA.4/5, BQ.1.1, and XBB.1.5 evaluated by FO-BLI biosensor at 35 (n= 9 mice per group, one experiment). Serum samples were diluted a hundred-fold against BA.4/5 and BQ.1.1, while a 10-fold against XBB.1.5 (C) Grouped serum neutralizing antibody activity against BA.4/5, BQ.1.1, and XBB.1.5. The serum from each mouse, which had been injected with multivalent or XBB.1.5 vaccine, was mixed into a single sample and assayed for neutralizing antibody activity (n= 9 mice per experiment).

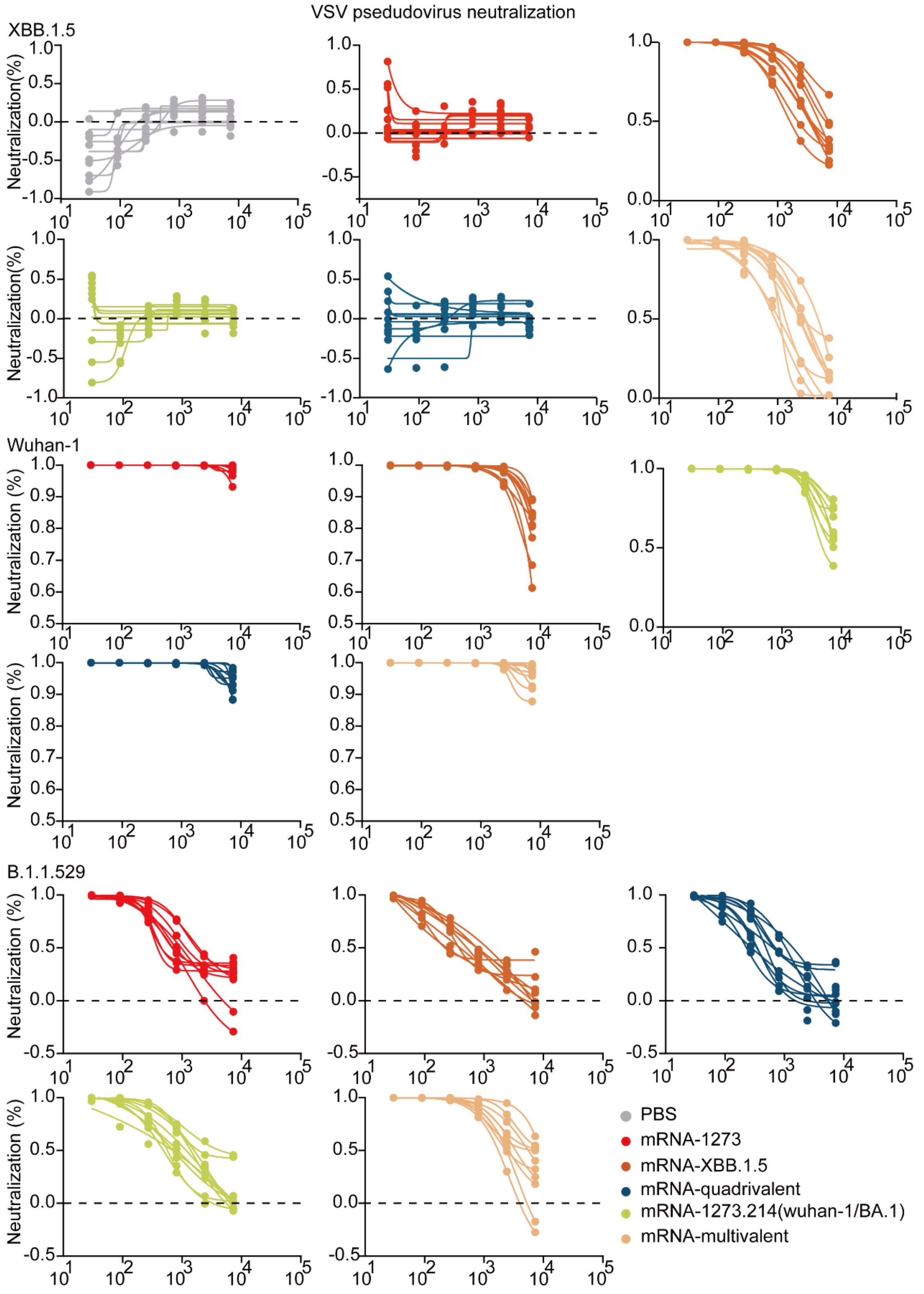


**Figure S3. Comparison of serum neutralization using VSV pseudoviruses expressing XBB.1.5, Wuhan-1, and B.1.1.529.** BALB/c mice were immunized with two 10 μg dose of mRNA-1273, mRNA-1273.214, mRNA-quadrivalent, mRNA-XBB.1.5, and mRNA-multivalent. (A) Serum neutralizing antibody activity at day 35 against XBB.1.5 assessed using VSV pseudoviruses. (B-C) Serum-neutralizing antibody activity at day 35 against wuhan-1 and B.1.1.529 assessed using VSV pseudoviruses. Representative neutralization curves corresponding to mice are shown for the indicated vaccines.

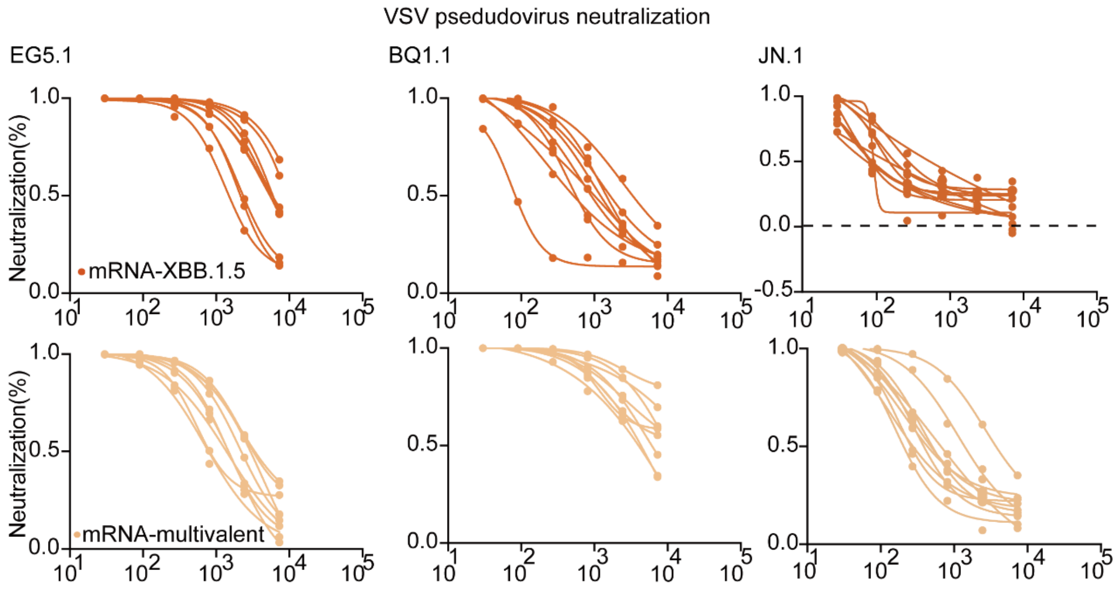


**Figure S4. Comparison of serum neutralization using VSV pseudoviruses expressing EG.5.1, BQ.1.1, and JN.1.** BALB/c mice were immunized with two 10 μg dose of mRNA-XBB.1.5 and mRNA-multivalent. Serum neutralizing antibody activity at day 35 against EG.5.1, BQ.1.1, and JN.1 assessed using VSV pseudoviruses.
